# A machine learning approach predicts essential genes and pharmacological targets in cancer

**DOI:** 10.1101/692277

**Authors:** Coryandar Gilvary, Neel S. Madhukar, Kaitlyn Gayvert, Miguel Foronda, Alexendar Perez, Christina S. Leslie, Lukas Dow, Gaurav Pandey, Olivier Elemento

## Abstract

Loss-of-function (LoF) screenings have the potential to reveal novel cancer-specific vulnerabilities, prioritize drug treatments, and inform precision medicine therapeutics. These screenings were traditionally done using shRNAs, but with the recent emergence of CRISPR technology there has been a shift in methodology. However, recent analyses have found large inconsistencies between CRISPR and shRNA essentiality results. Here, we examined the DepMap project, the largest cancer LoF effort undertaken to date, and find a lack of correlation between CRISPR and shRNA LoF results; we further characterized differences between genes found to be essential by either platform. We then introduce ECLIPSE, a machine learning approach, which combines genomic, cell line, and experimental design features to predict essential genes and platform specific essential genes in specific cancer cell lines. We applied ECLIPSE to known drug targets and found that our approach strongly differentiated drugs approved for cancer versus those that have not, and can thus be leveraged to identify potential cancer repurposing opportunities. Overall, ECLIPSE allows for a more comprehensive analysis of gene essentiality and drug development; which neither platform can achieve alone.

## INTRODUCTION

The identification of essential genes within cancer cells inform numerous therapeutic approaches and enable personalized medicine applications. For example, inhibiting the product of cancer specific essential genes can selectively kill cancer cells while presumably having no lethal effect on healthy cells^1^. In addition, drug treatments can be prioritized based on the essentiality of drug targets within specific cancer cells^2^. Cancer specific essential genes in conjunction with a patient’s unique mutational landscape can also be used to identify synthetic lethal relationships^3-6^. With such diverse applications in precision medicine, it has become increasingly important to accurately identify essential genes in a given sample.

One popular method for identifying cancer-specific essential genes is a genome-wide loss-of-function (LoF) screen in cancer cell lines. A related method, RNA interference (RNAi) has long been the standard method of conducting these screens and works by successfully suppressing expression of genes through targeting mRNA. Short hairpin RNA (shRNA) LoF screens have proven to be invaluable to cancer therapeutics through the identification of novel dependencies^1^ and target prioritization^7^. However, with the introduction of the CRISPR-Cas9 genome editing system, there has been a shift in many genome-wide LoF screens, creating a bias towards CRISPR over RNAi^8-10^. The CRISPR-Cas9 platform targets specific DNA sequences of genes, as opposed to the targeting of mRNA stability and translation by the RNAi machinery^11^, and disrupt genes by creating insertions and deletions (‘indels’) that in most cases lead to early termination or a frame shift within that gene, resulting in a knockout of gene expression.

While both CRISPR and RNAi can be used in the identification of essential genes, making the decision between platforms is often uninformed and leads to variable results^8, 9, 12^ due to the uncharacterized differences between technologies. In shRNA screens seed-mediated off-target effects can cause inaccurate essential gene call^13, 14^; however, even when screens account for these off-target effects and thus increase their accuracy, inefficient knockdown can lead to variable results within LoF screens^13^. In addition, numerous disadvantages to the CRISPR system have come to light,^15^ such as inaccurately calling genes with copy number amplifications as essential, or variability within screens arising from the creation of null alleles, heterozygous or wild type cells^16^. While both platforms suffer from clear weaknesses and are mechanistically distinct it is unsurprising that previous research has shown that CRISPR and shRNA LoF screens have low overlap in the essential genes they identify^12^.

Here, we characterize the genomic features of platform-specific essentiality and further identify systematic biases in both platforms. We demonstrate that the highest confidence essential genes arise when combining LoF results from both CRISPR and shRNA screens. Furthermore, we introduce ECLIPSE, a machine-learning platform that leverages genomic, cell line, and design features to predict context and platform-specific gene essentiality without any prior LoF screening data required. We demonstrate the value of such predictions by using ECLIPSE’s predictions to select efficacious drugs based on predicted gene essentiality. Our extensive characterization of platform specific essential genes and the development of ECLIPSE highlights the biological differences between platforms and allows for a more complete understanding of gene essentiality.

## RESULTS

### Significant discrepancies between shRNA and CRISPR screens

To evaluate how shRNA and CRISPR screens could be used together to find essential genes, we first turned to DepMap the largest, publically available shRNA and CRISPR LoF screening efforts across diverse cancer cell lines^15, 17^. For each gene-cell line pair, DepMap calculated an essentiality score – measuring the effect a given knockdown/knockout had on the survival of that specific cell line – with lower scores corresponding to more essential genes^18^. These scores were calculated using the DEMETER and CERES algorithm for shRNA and CRIPSR screens, respectively. The DEMETER algorithm specifically accounts for seed-mediate off-target effects^19^, while CERES corrects for the noted copy number bias within CRISPR screens. We found that when matching these gene-cell line pairs the correlation between shRNA and CRISPR scores lacked strong concordance (corr = 0.2, p <0.001,Spearman Correlation, **Fig S1A, Supplementary Data**). This low correlation could have drastic impacts as it indicates that certain essential genes could be either missed or incorrectly called based on the platform being used.

To decrease the inherent noise caused by genes that are not strongly essential (intermediate DEMETER and CERES scores), we next separated each gene-cell line pair into one of 4 categories based on the screening results from each platform: 1) called as essential only by shRNA, 2) only by CRISPR, 3) by both platforms, or 4) by neither (**Fig. 1A**). For each platform, the top depleted genes within each cell line were defined as essential (**Methods**). We observed that even for marked essential genes in shRNA there was a broad distribution of essentiality scores within CRISPR, and vice versa with a significant difference between the mean LoF scores (Mann-Whitney, p < 0.001, **Fig. 1B-C**). Platform specific essential (PSE) genes, genes called as essential exclusively by shRNA or CRISPR, were the majority of all essential genes. Therefore, even for supposed essential genes there is still a large degree of discordance between the platforms.

**Figure 1:**
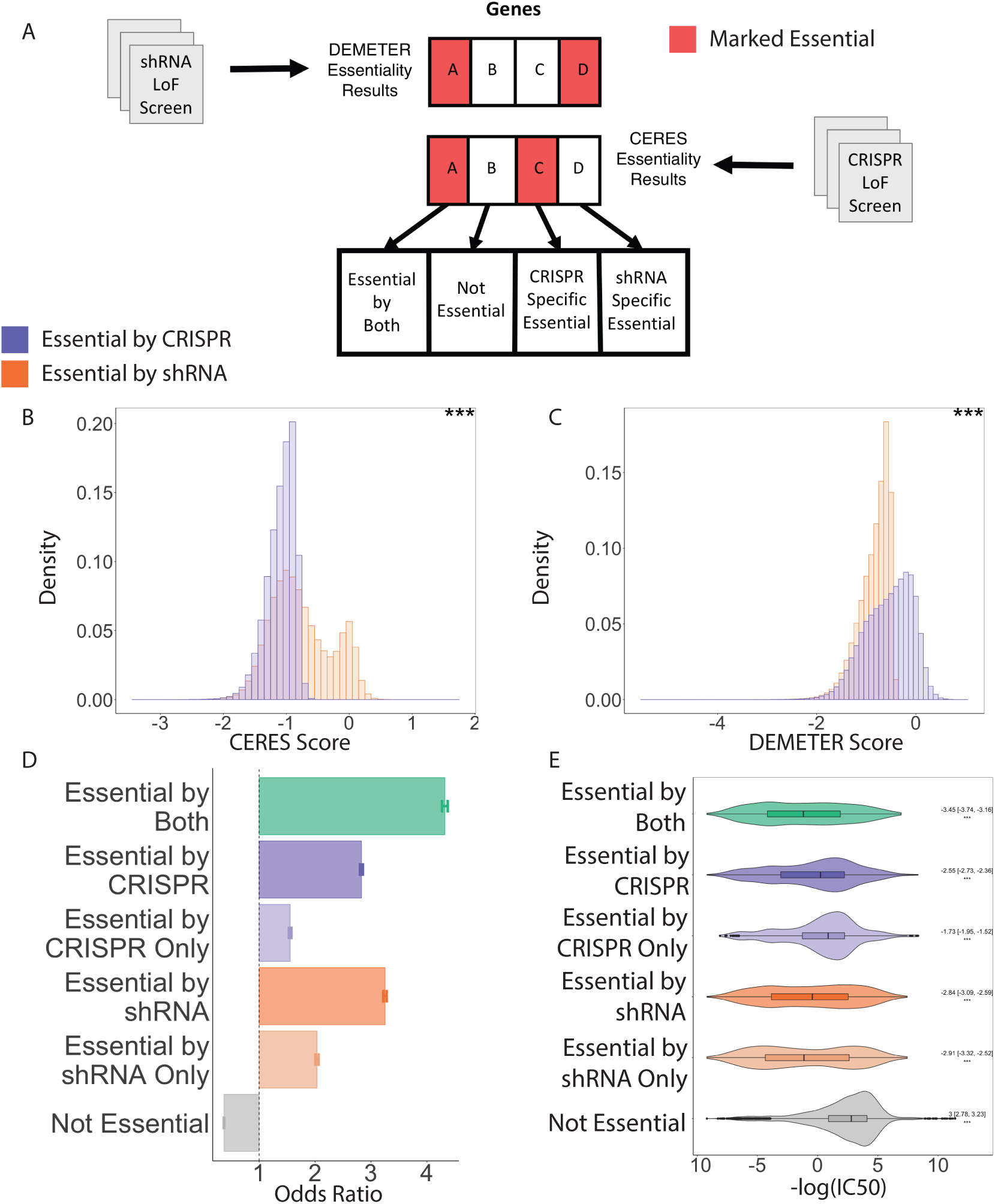
CRISPR and shRNA loss-of-function screens do not correlate. A) Schematic of the classification of gene essentiality within a cell line. B) Distributions of CRISPR CERES scores across two sets-genes classified as essential by CRISPR and genes classified essential by shRNA. C) Distributions of shRNA DEMETER scores across two sets-genes classified as essential by CRISPR and genes classified essential by shRNA. D) The odd’s ratio found by Fisher’s exact test of each category being true essential genes, as defined by loss-of-function intolerance and missense z-score, with each category having a significant p-value. E) The distribution of drug efficacy for drugs with at least one target marked essential by both or either category, location shift and p-value found using Mann-Whitney test.

We next evaluated both platforms in their ability to identify predetermined known essential genes. We first evaluated the performance of both shRNA and CRISPR in the categorization of the essential gene list found in Hart et al^20^, and found that both platforms performed extremely well (AUC = 0.97 for both, **Fig. S1B**). However, this set of essential genes was found using shRNA screens, therefore, to ensure a truly unbiased approach in the evaluation of each platform we identified a set of essential genes based on the frequency of loss of function and missense mutations across 60,706 individuals^21^. Additionally, we included the genes classified as “essential by both” to measure the performance of PSE and combined platform essentiality. To measure whether the discordance we observed was simply because one platform was drastically outperforming the other, we used this list as a “gold-standard” set of essential genes to objectively measure the accuracy of each platform (**Methods)**. We found that the essential genes marked by both platforms were more likely to be part of our gold-standard set (Fisher’s exact test, OR = 20.4, p <0.001) than genes marked as essential by any single platform (Fisher’s Exact Test, OR = 6.3, OR = 8.3, p <0.001, for shRNA and CRISPR, respectively) (**Fig. 1D, S2**). These findings indicate that the best identification of essential genes can be obtained only by combining the output of CRISPR and shRNA LoF screens.

In addition to the identification of essential genes, cancer LoF screenings can serve as a tool to identify effective drugs within certain cell lines. This is based on the hypothesis that if a gene is essential within a specific cell line, a specific inhibitor of that gene should efficiently kill cells grown from that cell line. To determine which type of essentiality best identified anti-cancer drugs, we obtained drug efficacy values for a set of 170 cell lines for 59 drugs, which had both shRNA and CRISPR screening data available for their targets (**Fig. 1E, Methods**). We found that there was a significant increase in a cell line’s sensitivity to a drug if at least one of that drug’s targets were marked as essential by both CRISPR and shRNA compared to drugs that had no targets marked as essential by both platforms (Mann-Whitney, Location Shift = −3.45, p <0.001). Interestingly, we observed platform specific differences in identifying effective drugs. For instance, we found that targets that were exclusively essential within shRNA screens had a larger increase in efficacy than targets that were exclusively essential within CRISPR (Mann-Whitney, Location shift = −2.91, p < 0.001 vs Location Shift = −1.73, p <0.001 for shRNA and CRISPR, respectively, **Fig, 1E**). This result indicates how shRNA screens may be more applicable for drug prioritization than their CRISPR counterparts. One possible explanation for this could be that the mechanism for shRNA knockdown is more similar to how some compound antagonism modes of action work than the DNA-targeted gene knockout induced by CRISPR^22^. However, the utilization of both CRISPR and shRNA, similar to with the identification of essential genes, is best when identifying effective drugs.

### Genomic features can help identify genes that will perform similarly across platforms

Diving deeper into the discordance observed between shRNA and CRISPR screens, we next sought to identify whether there were certain types of genes that produce similar results across the two platforms vs. those that give highly discordant scores. If we can identify which genes will give similar or opposing results across platforms, it could help determine the utility of performing both screening experiments rather than only a single one to test a single gene. For each gene, we measured the correlation between its essentiality scores for shRNA and CRISPR across all overlapping cell lines. We then separated genes into 3 categories based on this correlation: no correlation, negatively correlated (opposing results), positively correlated (similar results) (**Fig. 2A.**)

**Figure 2:**
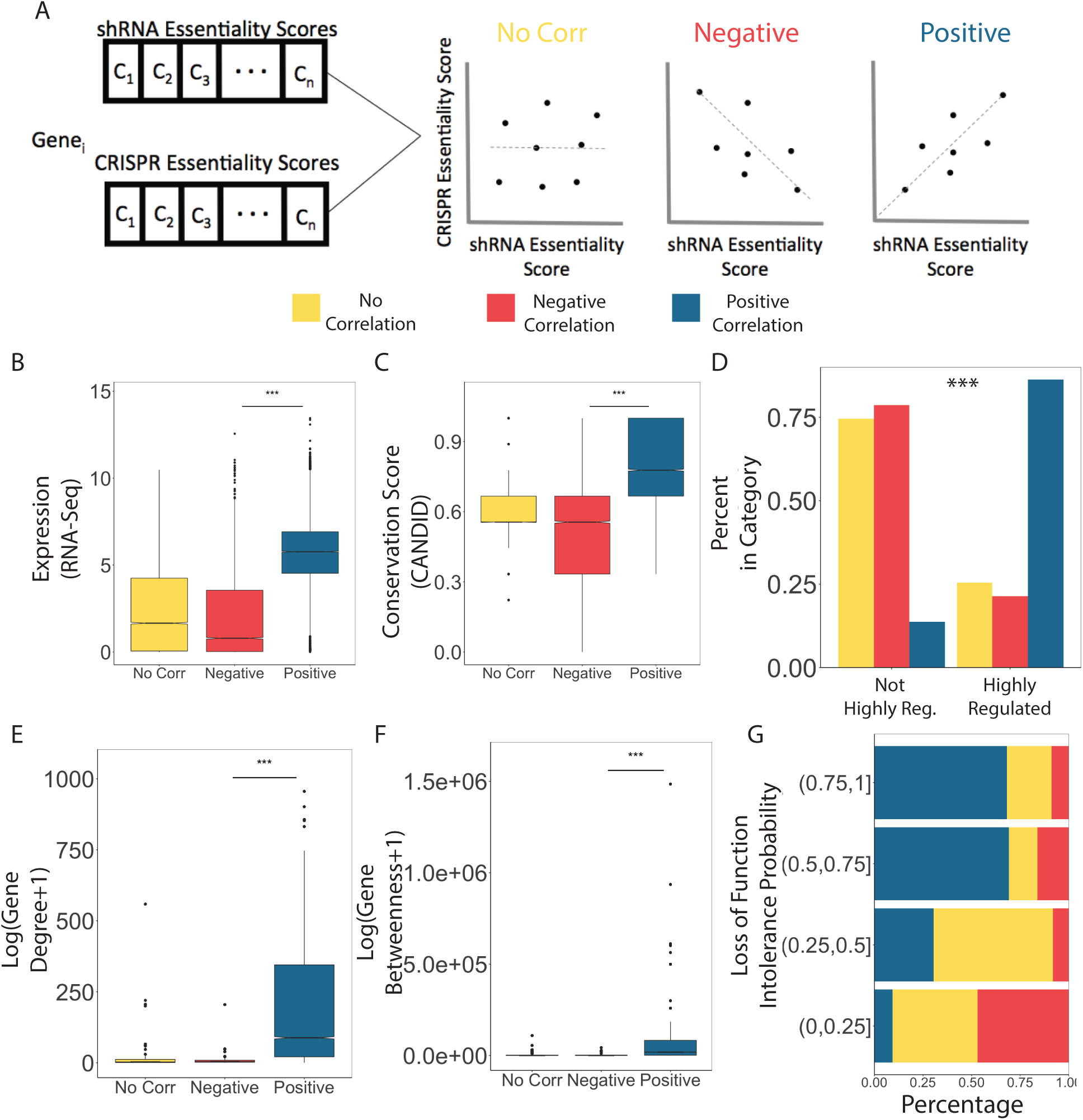
Genes that performed similarly between platforms share genomic attributes. A) Schematic of the classification of genes with positive, negative and no correlation between platforms across 16 cell lines. Genomic differences between genes of each correlation category with statistical significance found with the Kolmogorov-Smirnov test, for B) RNA-Seq gene expression, C) CANDID conservation scores D) the amount of transcription factor regulation (Fisher exact test), E) Gene Degree, F) Gene Betweenness and G) the distribution of each correlation type across differing LoF intolerance probabilities.

We found that many genomic features were able to accurately separate genes based on their cross-platform correlation (**Methods**). Genes that were highly correlated across the two platforms tended to be more highly expressed in that specific cell lines and were more highly conserved across species (**Fig. 2B-C**). In addition, we found that genes with similar results across platforms tended be more connected in a curated protein-protein interaction network (Methods, **Fig. 2E-F**). This pattern of highly connected genes performing similarly across platforms held true when examining the degree of regulation between genes and multiple transcription factors (**Fig. 2D**). Previous studies have shown that genes that are conserved, highly connected, and regulated by many transcription factors are often essential to survival^23, 24^. Therefore, we next compared cross-platform correlation for each gene to its measured intolerance to loss of function mutations^21^– the same feature we, and previous studies, have used to generate a gold standard list of essential genes. As expected, genes that were more intolerant to loss of function mutations (higher probabilities) were more likely to be positively correlated across platforms (**Fig. 2G**). This finding was promising as it indicates that, the comparison of results from both platforms will be consistent for this set of essential genes. However, we did observe that even at high LoF intolerance probabilities there were a significant number of genes that performed variably across the two platforms.

### Features related to Platform Specific Essentiality (PSE)

To further examine genes with discordant results, we focused on how numerous genomic and experimental design features differed across PSE genes, genes marked essential by both platforms and those that would be found essential using either platform. Focusing first on expression, we found that for the majority of the tested cell lines, genes identified as essential by both platforms were significantly more highly expressed than genes identified as essential by only one platform. Interestingly, PSE genes, genes exclusively marked essential by shRNA or CRISPR, also showed a difference in expression, with genes marked essential by CRISPR only have higher expression values (**Fig. 3A**). Previously essential genes have been found to have higher expression^23, 25^; therefore, the combined findings from both platforms appears to produce the highest confidence essential genes and CRISPR PSE genes also appear to follow more closely to classical essential gene characteristics. Examining other genomic features (**Methods**), we found a similar pattern where genes called as essential by both screens were more highly conserved and interconnected in biological networks than shRNA specific essential genes and CRISPR specific essential genes (**Fig. S3**). CRISPR PSE genes also demonstrated these characteristics as well, albeit at a much smaller effect size. These gene characteristics, high expression, high conservation and connected^23^, have been long known to be associated with evolutionarily essential genes. We tested if CRISPR was identifying cell line agnostic essential genes. To measure this, for each gene we calculated what percentage of the 379 cell lines each platform marked said gene as essential. We found that shRNA identifies significantly more cell line specific essential genes compared to CRISPR (**Fig. 3B**). Therefore, these differences in genomic features could point to the inherent variability between cancer specific essential genes being found within shRNA and not CRISPR.

**Figure 3:**
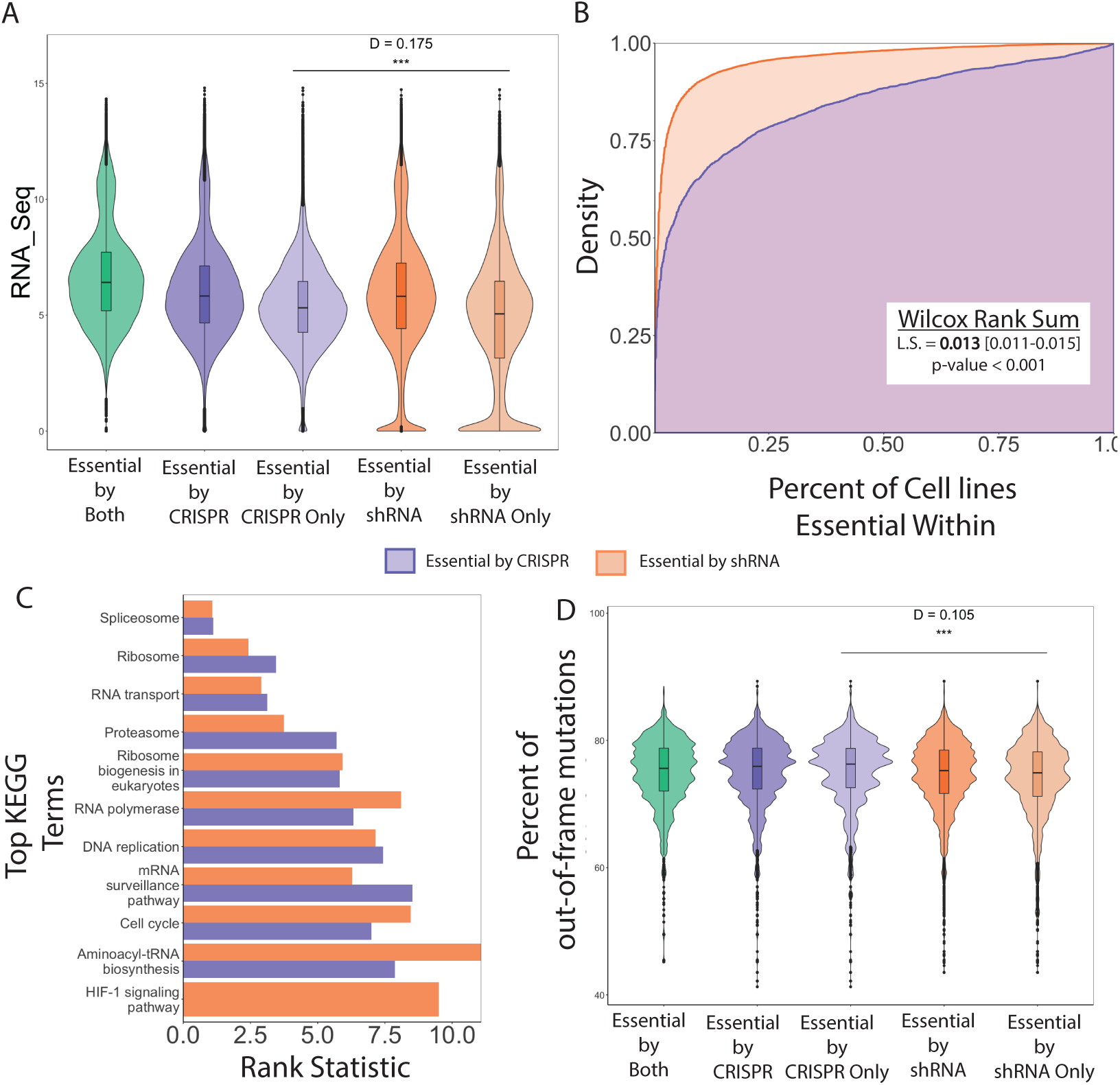
Genes marked essential by both platforms showed classic hallmarks of essential genes. A) The distribution of gene expression for PSE genes and all essential genes the Kolmogorov-Smirnov D statistic and p value are shown for the comparison of platform specific essential genes. B) The cumulative distribution of the percent of cell lines each gene is essential within for all genes found essential in shRNA or CRISPR LoF screenings. C) The top 10 KEGG pathway enrichment of genes marked essential by CRISPR (purple) and those marked essential by shRNA (orange), per each cell line, with the corresponding rank across all cell lines. D) The distribution of a predicted out of frame mutation, according to FORECasT, for all essential gene types.

Furthermore, certain differences in other genomic characteristics could point towards biases in either CRISPR or shRNA screens and these are important to consider when designing future screens. From our findings and previous work^12^, each platform identifies essential gene’s associated with unique pathways (**Fig. 3C**), such as shRNA essential genes being enriched for RNA polymerase, across all cell lines more consistently than CRISPR essential genes. While CRISPR essential genes are were consistently enriched for DNA replication more often than shRNA essential genes (**Methods**). Other platform specific biases, such as the design of the single-guide RNAs (sgRNA), a key component of the CRISPR-Cas9 system shown to influence knock-out efficiency^26-28^, proved to contribute to the discordance between shRNA and CRISPR. The likelihood of causing an out of frame mutation was significantly lower in shRNA specific essential genes compared to CRISPR specific essential genes (**Fig. 3D, S4**), which indicates erroneous results by CRISPR. Perhaps designing more specific sgRNAs for shRNA specific genes would have resulted in essential calls in CRISPR screens as well.

### ECLIPSE predicts Platform-Specific Essentiality

Based on these results we reasoned that combining shRNA and CRISPR screens could combat some of the pitfalls of each individual platform and provide more accurate essentiality screening results. However, due to costs – both in terms of time and money – it is often not feasible to use two separate technologies. To address this, we developed ECLIPSE – the Estimation of Combined LoF scores and Inference of Platform Specific Essentiality. ECLIPSE leverages the genomic features we found consistent with essential genes, such as gene expression, network properties and conservation, in combination with cell-line and design features to predict cell line specific essentiality results for both platforms, as well as results specific to either platform, without any prior LoF results (**Fig. 4A**).

**Figure 4:**
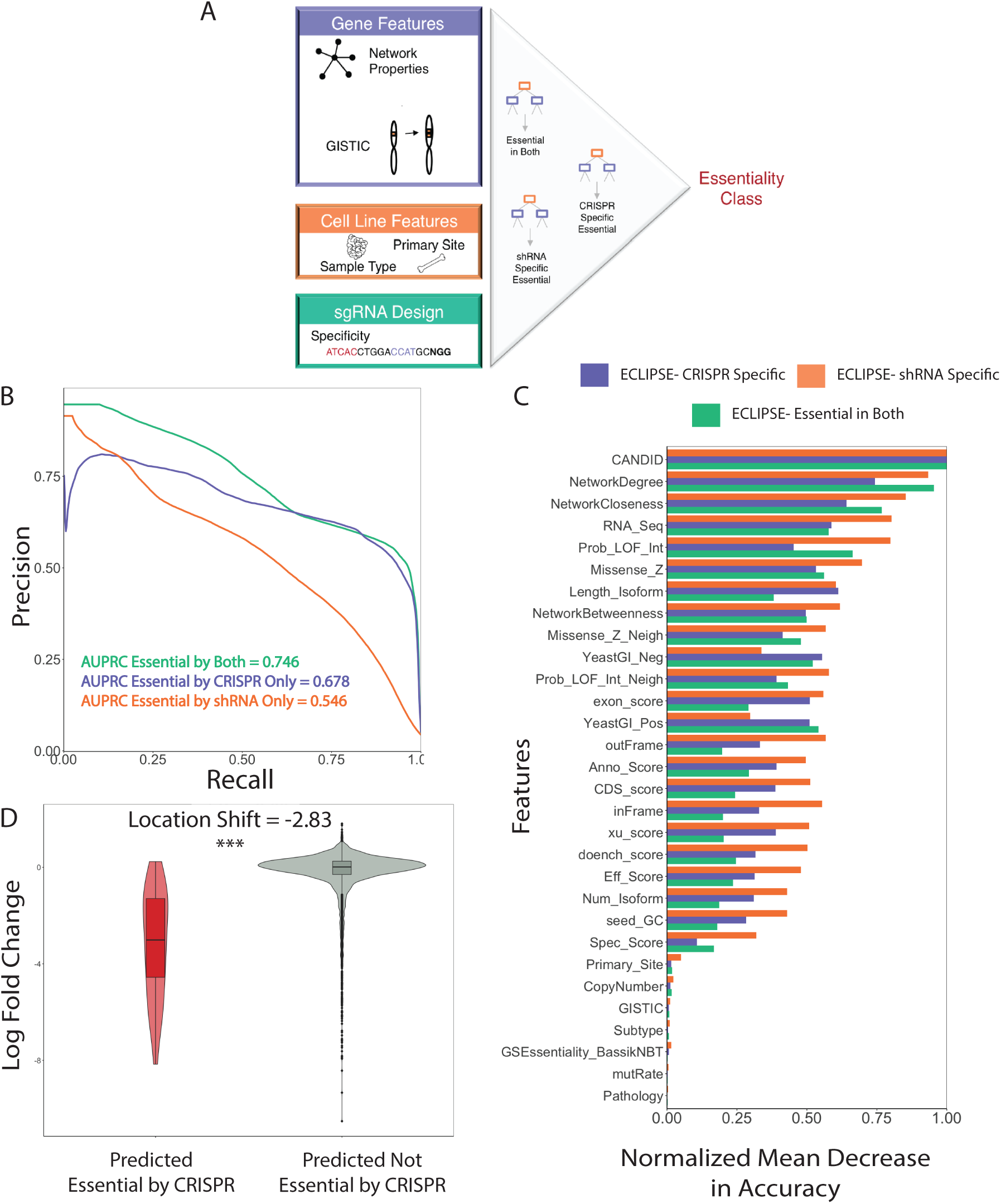
ECLIPSE can accurately predict high confidence and platform specific essential genes. A) Schematic of main data input to ECLIPSE, a random forest classification model. B) Precision-recall curve for ELCIPSE, with the classification of all three essentiality classes. C) The normalized feature importance of ECLIPSE for each classification class D) The distribution of the log fold change within the CRISPR test set for those predicted essential and not essential by ECLIPSE

For each essentiality type (i.e.: marked essential by both, CRISPR specific essential, and shRNA specific essential) a binary random forest classification model was built and 10-fold cross validation was used to evaluate performance (**Fig. 4A**). A separate model was built for each essentiality type classification due to the highly variable imbalance between classes (Table S1). For the same reason of class imbalance, we evaluated the performance of these models in terms of the class-specific Area Under the Precision-Recall Curve (AUPRC) measure^29^ in addition to the standard Area Under the Receiver Operating Characteristic (ROC) Curve (AUC) score^30^.

The identification of the highest confidence essential genes, those called as essential by both platforms, had significant predictive performance, achieving an AUC of 0.984 **(S5).** Due to the large class imbalance in all of these models the AUPRC expected by random is extremely low (AUPRC = 0.049), comparatively our model does significantly better and achieves an AUPRC of 0.746 (**Fig. 4B**).

To enable the prediction of PSE, features known to contribute to platform biases were included, such as sgRNA design and GC content of target strands. We found that ECLIPSE performed comparably on identifying platform specific essential genes, with significant predictive performance for shRNA specific essential and CRISPR specific essential genes (AUC = 0.981, AUC = 0.938, respectively, **Fig. S5**). Again, we analyzed the AUPRC and found that both PSE models performed better than random, with the CRISPR-specific and shRNA-specific essentiality models achieving AUPRC of 0.678 and 0.546 respectively, as compared to those of their random counterparts (0.049 and 0.044 respectively) (**Fig. 4B**).

To understand the features that contribute to each of ECLIPSE’s three classifications, we measured the mean decrease in accuracy upon removal of each feature (**Methods, Fig. 4C**). We found that features such as conservation, expression and network degree were important in each classification task. This analysis also revealed a subset of features that were differentially valuable based on the type of essentiality being predicted. For instance, we found that many of the sgRNA design features were more valuable in predicting shRNA specific essential genes than compared to any other classification. In addition, the number of isoforms of a gene was more informative in the classification of shRNA specific essential genes, which we hypothesize arises from the mechanistic differences between platforms. For example, shRNA is targeting mRNA, then if the seed sequence is not robust enough to target all isoforms, this could lead to incomplete knockdowns^31^.

To test the performance of our model we used the results from a CRISPR LoF screening conducted using kinase domain focused sgRNA library that has been previously published^32, 33^. In total, there were 11 cell lines tested that were not used within the training of ECLIPSE. We collected all available genomic, cell line and sgRNA design features as described above. We ran all gene-cell line pairs through both the CRISPR specific essential ECLIPSE model and essential by both ECLIPSE model. We found that the genes classified as essential by either ECLIPSE model had significantly lower log fold change compared to those predicted to be not essential (Mann-Whitney, Location Shift = −2.83, p < 0.001, **Figure 4C**). This CRISPR screen served as a valuable test set to prove the utility of ECLIPSE to predict LoF screens done using a diverse set of cell lines, genes and libraries.

### ECLIPSE accurately predicts pharmacological response

We next investigated whether ECLIPSE’s gene essentiality predictions in cell lines that did not have prior shRNA and CRISPR screening data are predictive of pharmacological response. We hypothesized that a drug will have better responses within cell lines where its target(s) is predicted to be more essential. Therefore, we evaluated 60 drugs that had publicly available pharmacological data across 212 cell lines. For each target-cell line pair we used ECLIPSE to predict the gene’s essentiality status, for all essentiality classes (**S6**). For targets predicted to be essential by both platforms, we found that drugs targeting these genes had significantly higher efficacy than drugs whose targets were predicted non-essential (Mann-Whitney, Location Shift = 2.6, p < 0.001). Additionally, drugs with gene targets predicted essential in shRNA only or CRISPR only were significantly more efficacious than those drugs with targets predicted not essential (Mann-Whitney, Location shift = 2.27 vs 2.31 for shRNA specific and CRISPR specific, respectively, p-value < 0.001).The ability to identify effective pharmacological response demonstrates that ECLIPSE can be applied to cell lines to identify possible therapies based on the cancer specific essentialities; which enables more precise drug selection and prioritization. Overall this highlights how ECLIPSE can be used not only to understand mechanisms of CRISPR and shRNA screens, but also to inform drug treatment in a cell-specific basis.

However, there are many situations where a drug can be highly efficacious in a cell line yet perform poorly in the clinic. For example, if a drug is targeting a gene that is essential within both cancerous and healthy cells, the drug might be highly toxic and likely not make it past Phase 1 clinical trials, if that. Therefore, we must consider two things, first that a gene will be essential within a cell line and second that a gene’s essentiality is highly specific (either to all cancer cells vs healthy cells or a specific cancer cell type). To do this we modified our predicted ECLIPSE score to create a Target Score, which is an integration of ECLIPSE scores across all cell lines (see **Methods**). The Target Score will equally weigh both efficacy and specificity. We next tested if our Target Score could be used to identify clinically relevant cancer therapeutics. We ran all known targets of 333 drugs approved to treat cancer and 6,013 drugs not currently approved or investigated to treat cancer through ECLIPSE and calculated the Target Score for each drug (**Methods**). We found that approved cancer drugs had significantly higher Target Scores compared to drugs not approved for any cancer indication (**Figure 5**). Overall, using the ECLIPSE essential by both model proved to be the most effective in the identification of cancer drugs (Mann-Whitney, Location Shift = 0.22, p-value < 0.001), however both CRISPR specific and shRNA specific models also showed statistical significance in the differentiation of cancer and non-cancer drugs. As expected, using only the raw ECLIPSE score in place of the Target Score did not show as great of a difference in approved and non-approved cancer drugs, however there was still a significant difference between the drug categories for each platform (**Fig. S7**). Overall, these results show a powerful opportunity for ECLIPSE to be applied to drug development in the identification of novel cancer targets and possible therapeutic repurposing.

**Figure 5:**
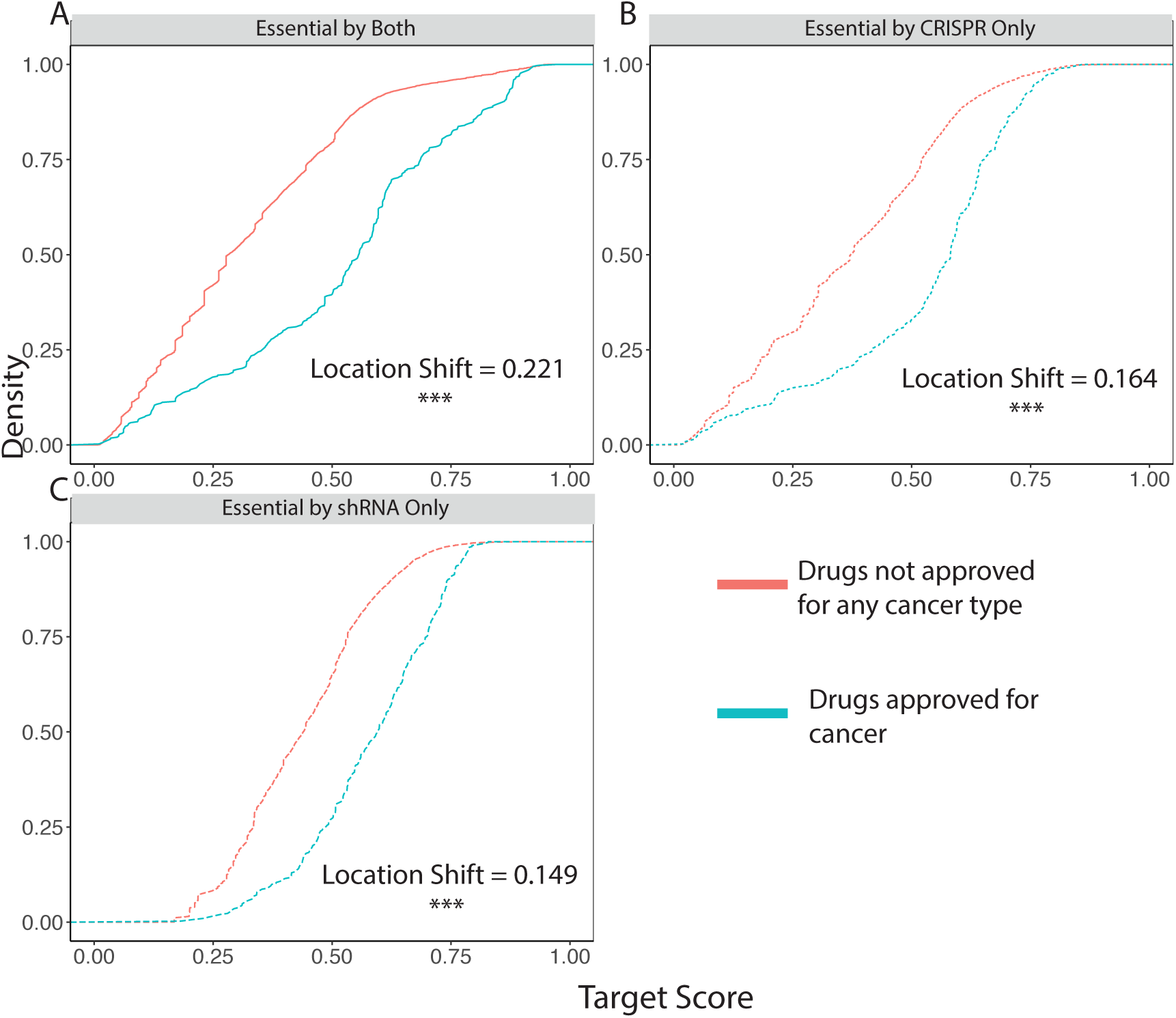
ECLIPSE Target Scores accurately distinguish approved cancer drugs. The cumulative distribution of the Target Scores for drugs approved for cancer and drugs not approved for cancer are shown, as calculated using A) ECLIPSE – essential in both B) ECLIPSE – essential in CRISPR only and C) ECLIPSE-essential in shRNA only

### ECLIPSE predicts new drug indications

Since ECLIPSE was able to accurately differentiate between approved cancer drugs and non-approved cancer drugs, we wanted to test if our ECLIPSE predictions could identify new cancer therapeutics based on predicted essentiality of their targets. We conducted these analyses using the previously calculated Target Scores based on the ECLIPSE predictions of essential in both, since that showed the highest statistical difference in identifying clinically relevant cancer drugs. We narrowed the list of 6,013 non-cancer drugs to exclude any drugs that shared a target with a drug approved or investigated to treat any cancer type (**Methods**). This ensured that the repurposing opportunities we identified using ECLIPSE would be novel. For each cancer type we ranked every drug based on the Target score, additionally for all drugs with DepMap screening information available for their targets we ranked all drugs by their CRISPR LoF screening score, shRNA LoF screening score and the maximum expression of that gene in that cancer type. Similar to our previous analysis we used the most extreme value of all cell lines for a primary site for each gene and then used the most extreme value of all drug targets, an extreme value was the maximum or minimum for expression data and LoF screening results, respectively (**Methods**).

Using our rankings, higher rank meaning the drug targets were more essential, we sought to evaluate if we could identify drug candidates which could not have been found using only either individual platform of LoF screenings or expression data. When specifically evaluating lung cancer, we found that Orilistat, a gastrointestinal lipase inhibitor (targeting genes: FASN, LIPF and PNLIP) currently approved for weight management, ranked the highest using the ECLIPSE rank. This drug also ranked second in skin cancer using the ECLIPSE score. Interestingly, previous preclinical work has shown that Orlistat decreases tumor cell proliferation and induced apoptosis within numerous prostate cancer^34^ and melanoma^35^ mouse studies, as well as initiating alternative energy pathways within non-small cancer cell lines^36^. Using ECLIPSE, Orlistat was easily flagged as a potential cancer therapeutic, however with traditional experimental or expression values Orlistat could have easily been overlooked (**Table 1**) – this example highlights the strength of applying ECLIPSE to drug repurposing.

**Table 1:** Novel cancer drug identification by ECLIPSE Target Score predictions. Drugs currently not approved or in clinical trials for cancer ranked highly as possible cancer therapeutics in numerous cancer types by ECLIPSE, compared with the rank found using CRISPR CERES, shRNA DEMETER scores or general expression.

| Drug | Current Indication | Cancer Type | ECLIPSE Rank | CRISPR Rank | shRNA Rank | Expression Rank |
| --- | --- | --- | --- | --- | --- | --- |
| Orlistat | Obesity | Lung | 1 | 49 | 65 | 45 |
|  |  | Skin | 2 | 24 | 79 | 43 |
| Memantine | Alzheimer's Disease | Bone | 6 | 138 | 107 | 319 |
|  |  | Breast | 24 | 144 | 73 | 332 |

In addition to orlistat, we identified memantine, an NMDA antagonist, as a promising new cancer treatment candidate for a subset of cancer types. Memantine was ranked 6^th^ for bone cancer and 24^th^ for breast cancer, using ECLIPSE predictions. Using either shRNA, CRISPR or expression memantine was ranked significantly lower for both cancer types (>100 for all techniques in bone cancer, **Table 1**). Previous research into NMDA antagonists have shown they can inhibit the extracellular signal regulated ½ kinase pathway and significantly decrease the proliferation of cancer cells^37^, adding support for this drug as an anti-cancer drug. Furthermore, memantine has previously been tested within breast cancer cell lines and showed a significant decrease in cancer cell growth^38^. Memantine was successfully found using ECLIPSE scores, however it would have been overlooked if we were only using shRNA and CRISPR due to the low rankings. Altogether, these results suggest that ECLIPSE shows great promise in repositioning non-cancer and cancer therapeutics for diverse other cancer types.

## DISCUSSION

Since the first application of RNAi, genome-wide loss-of-function (LoF) screens have been instrumental in the identification of essential genes, however as genetic LoF can have varying consequences and efficacies, the data from these screens needs to be thoughtfully examined. Previous reports have highlighted how RNAi platforms, such as shRNA, and the CRISPR-Cas9 system can have discordant results, however these analyses were done on only a single cell type without presenting any global feature based analysis. Here we compare shRNA and CRISPR screening results on a panel of 379 cancer cell lines spanning multiple different tumor types, which resulted in highly discordant results with many platform specific essential genes. These platform specific essential genes were enriched in diverse pathways and demonstrated unique genomic attributes. Interestingly we also found that shRNA screens are often more predictive of drug efficacy results than CRISPR screens, again stressing the important biological differences between platforms.

Based on these findings, we concluded that the combination of both screens led to better identification of “gold-standard” essential genes. We developed ECLIPSE, a machine learning model to predict cell line-specific PSE, that demonstrated high accuracy, sensitivity, and specificity, without the input of prior LoF screens. As seen within our results, the AUPRC was substantially lower than the AUC across all classifiers. This is due to the severe imbalance between essential and non-essential genes. Although AUC is routinely used as a performance metric in classification models, it can be misleading in cases of high-class imbalance, such as this one, due to AUC weighing classification errors in both classes equally. Since essentiality is a rarity amongst protein-coding genes, our reported AUPRCs are more appropriate when measuring the predictive power of ECLISPE, since AUPRC enables us to assess the classifiers’ performance on the minority classes. For all four of our models, the AUPRC is significantly higher than what would be observed at random and give promising predictions regarding a gene’s essentiality status.

ECLIPSE predictions not only provide information on the essentiality status of genes, but also incorporate the PSE, allowing a deeper understanding of the type of essentiality and the pitfalls – such as CRISPR sgRNA design – in each platform. ECLIPSE can be used with CRISPR screens to mark genes for which CRISPR results may need extra validation (CRISPR-specific genes). Finally, we demonstrated ECLIPSE’s utility in treatment prioritization as using ECLIPSE’s predictions for drug targets led to the identification of more efficacious drugs for a specific sample.

While ECLIPSE shows significant predictive power in identifying genes that will be called as essential by both platforms and platform specific essential genes, we are limited by inherent noise within shRNA screenings. While in recent work there has been an overwhelming number of CRISPR design tools available, the analytics of shRNA screenings are being overlooked. We found using the LoF screening results that shRNA outperformed CRISPR in drug prioritization and believe this was not captured in the predicted results due to the uncharacterized noise within shRNA experiments. As additional work is done to evaluate the efficacy and accuracy of shRNA screens, we believe this data can easily be incorporated into our model to improve upon this work. Additionally, as more optimized libraries for CRISPR^39^ are performed in large scale screenings we believe our reliance on CRISPR analytic tools will decrease.

Our approach has the potential to inform experimentation, while simultaneously enhancing the understanding of gene essentiality. ECLIPSE can be used, prior to performing shRNA or CRISPR on a set of genes, for better hit prioritization. In addition, the success of ECLIPSE in diverse cell lines allows it to be used to find candidate essential genes on any cell line of interest. Prioritization of drug treatment in specific primary sites can also be informed by the cell-line specific essentiality profile. Overall our results characterize the effects of mechanistic differences between shRNA and CRISPR, while providing a predictive model, ECLIPSE, which can be used to better understand gene essentiality, which can lead to real clinical impact.

## METHODS

### Genome-wide Loss-of-Function Data

The loss-of-function essentiality data was downloaded via DepMap. The shRNA and CRISPR LoF screening results were collected from DepMap 18Q4^13, 40^. Pre-calculated DEMETER and CERES values were used to determine essentiality. Scores were then paired together for each gene in each specific cell line cell line creating ∼4.5 million cell line-gene pairs. Essential genes for each cell line were determined with a top 5% essentiality score cut off. For each gene the percent of the total cell lines of the 379 that a gene was marked essential within for each platform was found.

### Feature Collection

#### Genomic Features

For the 14830 genes tested on each platform multiple gene features were collected. Cell line specific mRNA expression and DNA copy number data was downloaded from the Cancer Cell Line Encyclopedia (CCLE). GISTIC 2.0 was used to calculate copy numbers through GenePattern. Conservation scores were found by using web-based tool CANDID^41^. The ExAC database provided mutational prevalence data, missense z score and the probability of loss-of-function intolerance^21^. The transcription-factor regulation status of a gene was found by downloading from ENCODE. Genes being regulate by 2/3 or more TFs were classified as being highly regulated.

#### Network Features

We curated a biological network that contains 22,399 protein-coding genes, 6,679 drugs, and 170 TFs. The protein-protein interactions represent established interaction^42-44^, which include both physical (protein-protein) and non-physical (phosphorylation, metabolic, signaling, and regulatory) interactions. The drug-protein interactions were curated from several drug target databases^44^.

#### sgRNA Features

The sgRNAs used within the genome-wide CRISPR screens that contributed to each gene’s essentiality score was found and downloaded via Project Achilles. E-CRISPR, a web based tool, was run to determine Xu-score, exon score, and GC content^26, 45, 46^_.._ Additionally, the FORECasT algorithm was used to predict if sgRNAs will cause an^47^ out of frame mutation.

### KEGG Pathway Analysis

KEGG pathway over representation was found using Limma R package^48^ for all genes marked as essential by shRNA and CRISPR. For each cell line the pathways that were significantly over represented were found and ranked for each platform. Then the rank statistic (the sum of the ranks of the length of cell lines) was found for each platform.

### Cross Platform Correlation

Across all cell lines the correlation between CRISPR LoF scores and shRNA LoF scores were found for each gene. The genes with the top 100 and bottom 100 Pearson coefficients were classified as being positively correlated and negatively correlated, respectively. The genes with the middle 100 Pearson coefficients were classified as having no correlation. For RPKM expression, CANDID conservation scores, gene degree and gene betweenness a Kolmogorov-Smirnov test was performed between the negative and positive correlated genes. The enrichment of highly regulated genes was found using a Fisher’s exact test.

### The ECLIPSE Model

ECLIPSE was trained using the features described above on the DepMap data of 379 overlapping cell lines. Random forest, a decision tree model, was used after model selection and implemented using the R statistical software with the randomForest package^49^. To evaluate predictive power 10-fold cross validation was used. For the binary models predicting essentiality class, down sampling was the chosen sub-sampling approach applied to each fold to account for class imbalances.

### Classification Evaluation

For evaluating all the binary essentiality classifications, receiver operating characteristic (ROC) and precision-recall curve (PRC) curves were created in R using the pROC^50^ and precrec^51^ packages respectively. Area-under-the-ROC curve (AUC) and area-under-the-PRC (AUPRC) scores were used to evaluate model performance.

### Drug Pharmacological Inhibition Analysis

Drug efficacy values for 60 drugs and 374 cell lines were downloaded from The Genomics of Drug Sensitivity in Cancer Project^52^. All drug targets were obtained from DrugBank Version 5.0 with a custom python web scrapping script^53^.

### Known Drug Efficacy

Drugs were narrowed down to those with targets that had both shRNA and CRISPR essentiality values available. 170 cell lines were found to have drug sensitivity data available for these drugs. A Kolmogorov-Smirnov test was performed between drugs with at least one target found to be essential, according to different essentiality classifications, and those that were not.

### Predicted Drug Target Essentiality

In total, we tested 60 drugs with known targets and drug sensitivity data available in 374 cell lines, cell lines used to train the model were excluded. For each target ECLIPSE was run to obtain essentiality predictions within each of the 213 cell lines. For each group a Kolmogorov-Smirnov test was performed to determine the cell line sensitivity difference between drugs with a least one target predicted to be essential in that cell line and those with none, for each essentiality class.

### Approved Anti-Cancer Drug Analysis

All approved drug indications were obtained from DrugBank Version 5.0 with a custom python web scrapping script^53^. In addition, clinical trial information was collected from AACT, a relational database with all clinicaltrial.gov information, using a custom R script.

We evaluated 6,013 drugs with available indication and target data, 333 being approved or within clinical trials for neoplastic diseases that we could accurately match to a cancer type included in our training set. For each drug the representative ECLIPSE score was the maximum ECLIPSE score of all targets of the maximum score of those targets in all cell lines of the matching primary site. A target score was also calculated which was a modification of the ECLIPSE score. First each gene was given a cell line normalized score, which was all the predicted scores for that gene across all cell lines normalized from 0-1. The average of that normalized score and the predicted score was then used as the Target Score. The maximum Target Score across all drug targets in all cell lines of a given primary site was used for each drug, similar to ECLIPSE score. A Mann-Whitney test was used to find the difference in predicted scores between drugs approved for cancer treatment versus drugs that are not approved or investigated for cancer treatment.

### Novel Drug Indication Analysis

To predict possible repurposing opportunities, we used the list of 6,013 non-cancer drugs. This was refined to include only approved drugs and those that did not share any molecular targets with approved cancer drugs. We then ranked drugs based on their target score, CRISPR LoF screening results^40^, shRNA LoF screening results^13^ or gene expression. For LoF screenings the minimum score of all known targets across all cell lines of the appropriate primary site was used. For expression data the maximum RPKM, collected from CCLE^54^, of all targets across all cell lines of the matching primary site was used. The comparison of ranks was then used to identify novel cancer therapeutics.

### Code Accessibility

Code is available upon request.

## ACKNOWLEDGEMENTS

The authors would like to thank the J. Mezey and the Elemento laboratory members for their feedback and discussions.

## AUTHOR CONTRIBUTIONS

C.M.G., N.S.M., and O.E. conceived, designed and developed the methodology for this work. C.M.G., N.S.M., K.M.G, and O.E. analyzed and interpreted the data. C.M.G. executed the machine learning analyses and wrote the initial draft of the manuscript. M.F. and L.D. provided CRISPR screening data and guidance. G.P. provided machine learning guidance. A.P. and C.L. executed the CRISPR analyses to extract additional parameters. O.E. supervised the study. All the authors reviewed and approved the manuscript.

## Supplementary Figures

Figure S1: CRISPR and shRNA loss-of-function screens do not correlate. A) Density plot showing the correlation between CRISPR ATARiS scores and shRNA ATARiS scores for gene-cell line pairs. The darker area corresponds to a higher density of values and correlation and R^2^ values were calculated using Spearman correlation (p < 0.001). B) The ROC plot of both CRISPR and shRNA scores in identifying Hart et al essential genes.

Figure S2: Genes marked essential by both platforms show essentiality characteristics. Violin plots for showing the range of A) loss-of-function intolerance probability and B) missense z-scores for genes in each essentiality class. C) The ROC plot of both CRISPR and shRNA scores in identifying the ExAC gold standard essential genes.

Figure S3: Classic essentiality characteristics vary for each essentiality class. Range of A) network degree, B) network betweenness and C) CANDID conservation scores, for platform specific essential genes and genes marked essential by both platforms, statistical significance found by Kolmogorov-Smirnov test.

Figure S4: CRISPR sgRNA design characteristics vary for each essentiality class. Range of A) Xu Score, B) Doench Score, C) Exon Score and C) GC seed score, for platform specific essential genes and genes marked essential by both platforms, statistical significance found by Kolmogorov-Smirnov test.

Figure S5: ECLIPSE model performs well in cross validation. A) The ROC of each ECLIPSE model with the associated AUC. B) The score for various model evaluation measures for each ECLIPSE model.

Figure S6: ECLIPSE scores differentiate efficacious drugs. A) The cumulative distribution graph of the drug efficacy of drugs with at least one target predicted essential by each ECLIPSE model vs those that did not.

Figure S7: ECLIPSE scores in differentiating approved cancer drugs and other drugs. The cumulative distribution of ECLIPSE scores for approved cancer drugs and other drugs for A) essential in both ECLIPSE model, B) PSE CRIPSR ECLIPSE model and C) PSE shRNA ECLIPSE. Statistical significance and location shift found by Mann-Whitney.

## Supplementary Data

File 1: The correlations between shRNA DEMETER and CRISPR CERES scores for each individual cell line. Both Spearman and Pearson Correlation statistics and p-values are given.

## References

1. Luo, B. et al. Highly parallel identification of essential genes in cancer cells. Proceedings of the National Academy of Sciences 105, 20380–20385 (2008).

2. Guidi, A. et al. Application of RNAi to genomic drug target validation in schistosomes. PLoS neglected tropical diseases 9, e0003801 (2015).

3. Jerby-Arnon, L. et al. Predicting cancer-specific vulnerability via data-driven detection of synthetic lethality. Cell 158, 1199 (2014).

4. Luo, J. et al. A genome-wide RNAi screen identifies multiple synthetic lethal interactions with the Ras oncogene. Cell 137, 835–848 (2009).

5. Hoffman, G.R. et al. Functional epigenetics approach identifies BRM/SMARCA2 as a critical synthetic lethal target in BRG1-deficient cancers. Proceedings of the National Academy of Sciences of the United States of America 111, 3128 (2014).

6. Madhukar, N.S., Elemento, O. & Pandey, G. Prediction of genetic interactions using machine learning and network properties. Frontiers in bioengineering and biotechnology 3, 172 (2015).

7. Rao, D.D., Vorhies, J.S., Senzer, N. & Nemunaitis, J. siRNA vs. shRNA: similarities and differences. Advanced drug delivery reviews 61, 746–759 (2009).

8. Boettcher, M. & McManus, M.T. Choosing the right tool for the job: RNAi, TALEN, or CRISPR. Molecular cell 58, 575 (2015).

9. Evers, B. et al. CRISPR knockout screening outperforms shRNA and CRISPRi in identifying essential genes. Nature biotechnology 34, 631–635 (2016).

10. Wang, T., Wei, J.J., Sabatini, D.M. & Lander, E.S. Genetic screens in human cells using the CRISPR-Cas9 system. Science (New York, N.Y.) 343, 80 (2014).

11. Boettcher, M. & McManus, M.T. Choosing the right tool for the job: RNAi, TALEN, or CRISPR. Molecular cell 58, 575–585 (2015).

12. Morgens, D.W., Deans, R.M., Li, A. & Bassik, M.C. Systematic comparison of CRISPR/Cas9 and RNAi screens for essential genes. Nature biotechnology 34, 634 (2016).

13. Jaiswal, A. et al. Seed-effect modeling improves the consistency of genome-wide loss-of-function screens and identifies synthetic lethal vulnerabilities in cancer cells. Genome medicine 9, 51 (2017).

14. Jackson, A.L. & Linsley, P.S. Recognizing and avoiding siRNA off-target effects for target identification and therapeutic application. Nature reviews. Drug discovery 9, 57 (2010).

15. Aguirre, A.J. et al. Genomic Copy Number Dictates a Gene-Independent Cell Response to CRISPR/Cas9 Targeting. Cancer discovery 6, 914 (2016).

16. Shalem, O., Sanjana, N.E. & Zhang, F. High-throughput functional genomics using CRISPR-Cas9. Nature reviews. Genetics 16, 299 (2015).

17. Cowley, G.S. et al. Parallel genome-scale loss of function screens in 216 cancer cell lines for the identification of context-specific genetic dependencies. Scientific data 1 (2014).

18. Shao, D.D. et al. ATARiS: computational quantification of gene suppression phenotypes from multisample RNAi screens. Genome research 23, 665 (2013).

19. Tsherniak, A. et al. Defining a cancer dependency map. Cell 170, 564–576. e516 (2017).

20. Hart, T., Brown, K.R., Sircoulomb, F., Rottapel, R. & Moffat, J. Measuring error rates in genomic perturbation screens: gold standards for human functional genomics. Molecular systems biology 10, 733 (2014).

21. Lek, M. et al. Analysis of protein-coding genetic variation in 60,706 humans. BioRxiv, 030338 (2016).

22. Housden, B.E. et al. Loss-of-function genetic tools for animal models: cross-species and cross-platform differences. Nature Reviews Genetics 18, 24–40 (2017).

23. Yang, L. et al. Analysis and identification of essential genes in humans using topological properties and biological information. Gene 551, 138–151 (2014).

24. Wang, T. et al. Identification and characterization of essential genes in the human genome. Science 350, 1096–1101 (2015).

25. Marcotte, R. et al. Essential gene profiles in breast, pancreatic, and ovarian cancer cells. Cancer discovery 2, 172–189 (2012).

26. Heigwer, F., Kerr, G. & Boutros, M. E-CRISP: fast CRISPR target site identification. Nature methods 11, 122–123 (2014).

27. Perez, A.R. et al. GuideScan software for improved single and paired CRISPR guide RNA design. Nature biotechnology 35, 347 (2017).

28. Zhang, X.-H., Tee, L.Y., Wang, X.-G., Huang, Q.-S. & Yang, S.-H. Off-target effects in CRISPR/Cas9-mediated genome engineering. Molecular Therapy— Nucleic Acids 4, e264 (2015).

29. Lever, J., Krzywinski, M. & Altman, N. (Nature Publishing Group, 2016).

30. Saito, T. & Rehmsmeier, M. The precision-recall plot is more informative than the ROC plot when evaluating binary classifiers on imbalanced datasets. PloS one 10, e0118432 (2015).

31. Kisielow, M., Kleiner, S., Nagasawa, M., Faisal, A. & Nagamine, Y. Isoform-specific knockdown and expression of adaptor protein ShcA using small interfering RNA. Biochemical Journal 363, 1–5 (2002).

32. Foronda, M. et al. Tankyrase inhibition sensitizes cells to CDK4 blockade. bioRxiv, 677823 (2019).

33. Tarumoto, Y. et al. LKB1, salt-inducible kinases, and MEF2C are linked dependencies in acute myeloid leukemia. Molecular cell 69, 1017–1027. e1016 (2018).

34. Kridel, S.J., Axelrod, F., Rozenkrantz, N. & Smith, J.W. Orlistat is a novel inhibitor of fatty acid synthase with antitumor activity. Cancer research 64, 2070–2075 (2004).

35. Carvalho, M.A. et al. Fatty acid synthase inhibition with Orlistat promotes apoptosis and reduces cell growth and lymph node metastasis in a mouse melanoma model. International journal of cancer 123, 2557–2565 (2008).

36. Sankaranarayanapillai, M., Zhang, N., Baggerly, K.A. & Gelovani, J.G. Metabolic shifts induced by fatty acid synthase inhibitor orlistat in non-small cell lung carcinoma cells provide novel pharmacodynamic biomarkers for positron emission tomography and magnetic resonance spectroscopy. Molecular Imaging and Biology 15, 136–147 (2013).

37. Stepulak, A. et al. NMDA antagonist inhibits the extracellular signal-regulated kinase pathway and suppresses cancer growth. Proceedings of the National Academy of Sciences 102, 15605–15610 (2005).

38. North, W.G., Gao, G., Memoli, V.A., Pang, R.H. & Lynch, L. Breast cancer expresses functional NMDA receptors. Breast cancer research and treatment 122, 307–314 (2010).

39. Doench, J.G. et al. Optimized sgRNA design to maximize activity and minimize off-target effects of CRISPR-Cas9. Nature biotechnology 34, 184 (2016).

40. Meyers, R.M. et al. Computational correction of copy number effect improves specificity of CRISPR–Cas9 essentiality screens in cancer cells. Nature genetics 49, 1779 (2017).

41. Hutz, J.E., Kraja, A.T., McLeod, H.L. & Province, M.A. CANDID: a flexible method for prioritizing candidate genes for complex human traits. Genetic epidemiology 32, 779 (2008).

42. Das, J. & Yu, H. HINT: High-quality protein interactomes and their applications in understanding human disease. BMC systems biology 6, 92 (2012).

43. Khurana, E., Fu, Y., Chen, J. & Gerstein, M. Interpretation of genomic variants using a unified biological network approach. PLoS computational biology 9, e1002886 (2013).

44. Aksoy, B.A. et al. PiHelper: an open source framework for drug-target and antibody-target data. Bioinformatics 29, 2071–2072 (2013).

45. Xu, H. et al. Sequence determinants of improved CRISPR sgRNA design. Genome research 25, 1147 (2015).

46. Doench, J.G. et al. Rational design of highly active sgRNAs for CRISPR-Cas9-mediated gene inactivation. Nature biotechnology 32, 1262 (2014).

47. Allen, F. et al. Predicting the mutations generated by repair of Cas9-induced double-strand breaks. Nature biotechnology 37, 64 (2019).

48. Ritchie, M.E. et al. limma powers differential expression analyses for RNA-sequencing and microarray studies. Nucleic acids research 43, e47–e47 (2015).

49. Liaw, A. & Wiener, M. Classification and regression by randomForest. R news 2, 18–22 (2002).

50. Robin, X. et al. pROC: an open-source package for R and S+ to analyze and compare ROC curves. BMC bioinformatics 12, 77 (2011).

51. Saito, T. & Rehmsmeier, M. Precrec: fast and accurate precision–recall and ROC curve calculations in R. Bioinformatics 33, 145–147 (2017).

52. Yang, W. et al. Genomics of Drug Sensitivity in Cancer (GDSC): a resource for therapeutic biomarker discovery in cancer cells. Nucleic acids research 41, D955–D961 (2012).

53. Law, V. et al. DrugBank 4.0: shedding new light on drug metabolism. Nucleic acids research 42, D1091–D1097 (2013).

54. Barretina, J. et al. The Cancer Cell Line Encyclopedia enables predictive modelling of anticancer drug sensitivity. Nature 483, 603 (2012).

