## Supplementary figures and images for "A machine learning approach predicts essential genes and pharmacological targets in cancer"

### Supplemental Figures

A

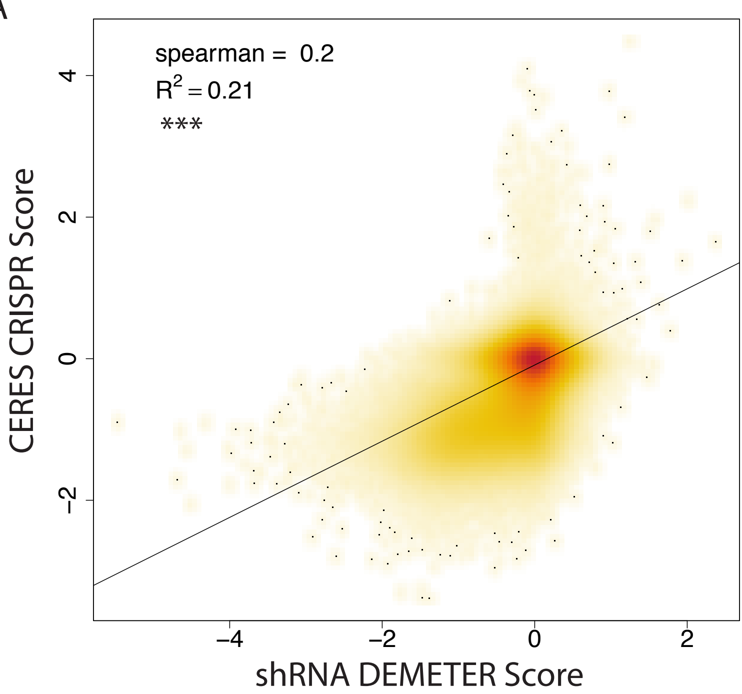


#### ROC – P: 63026, N: 214334

B

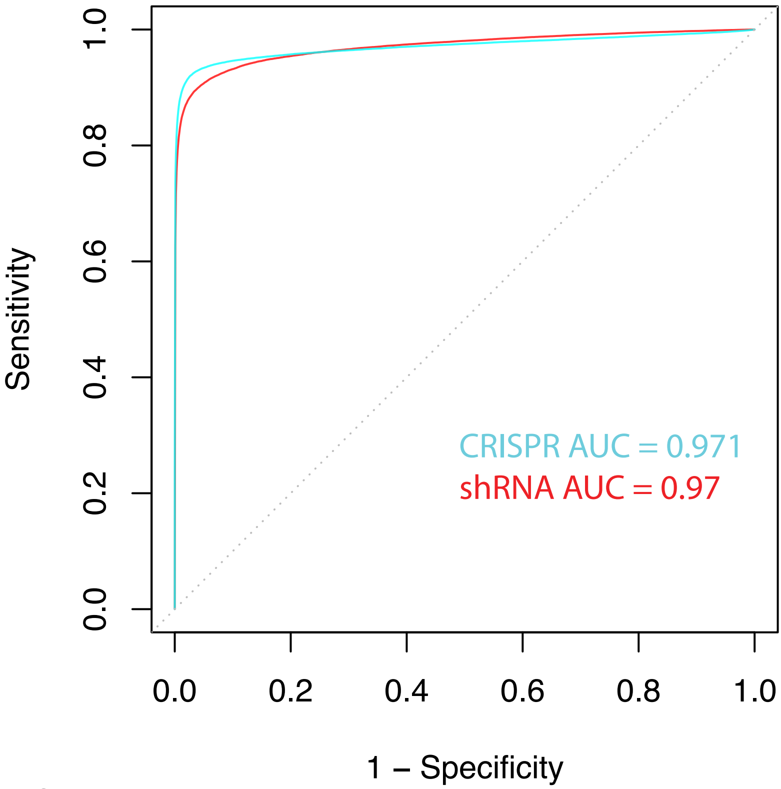


Figure S1

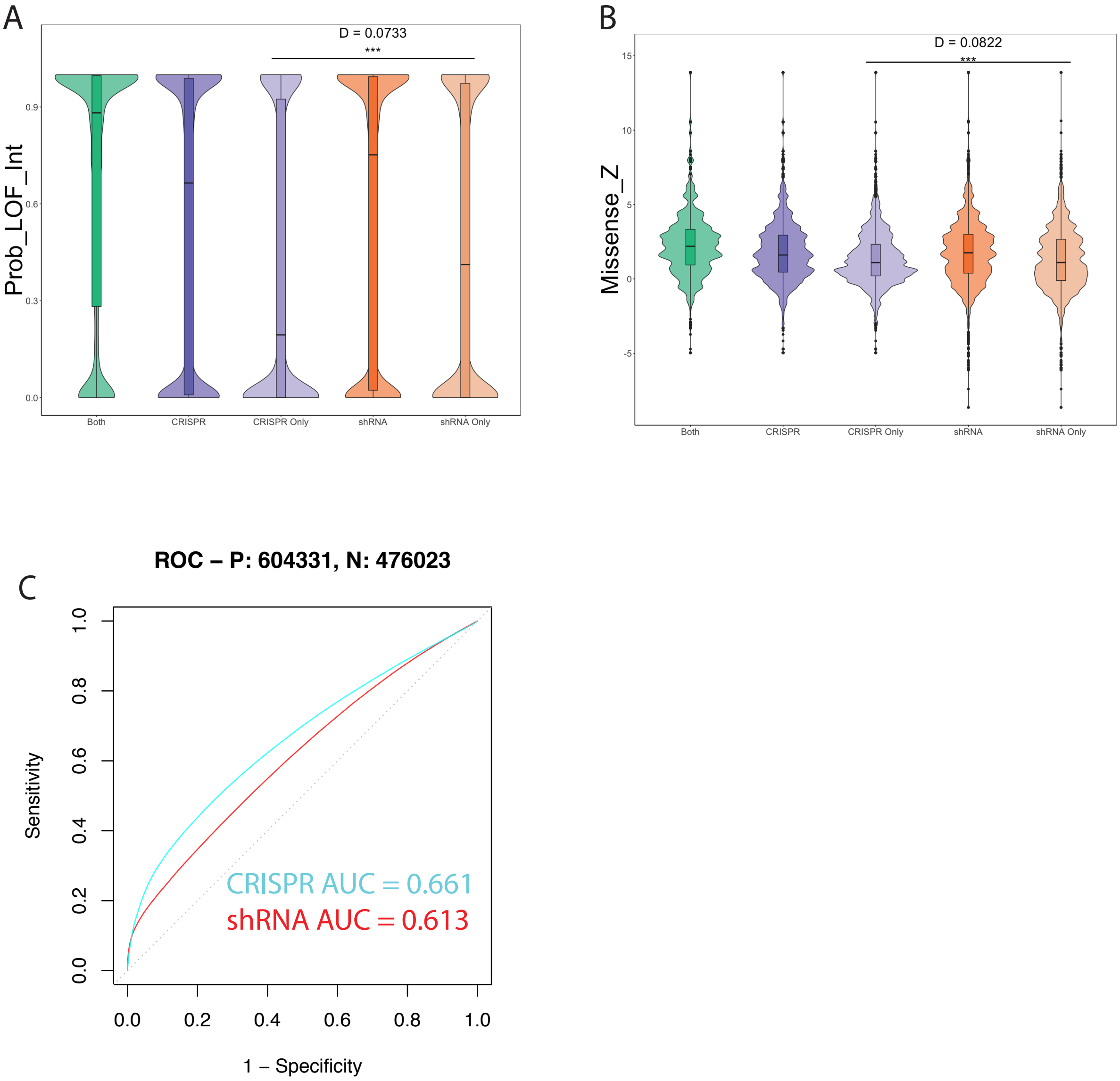

A

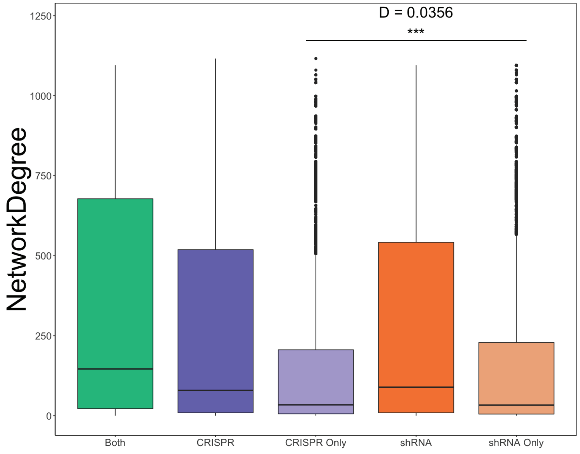


B

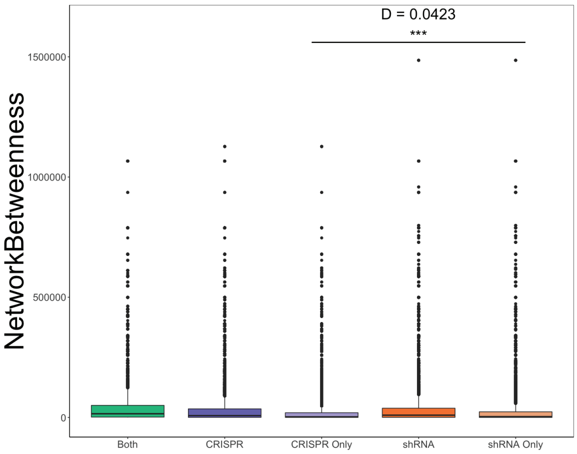


C

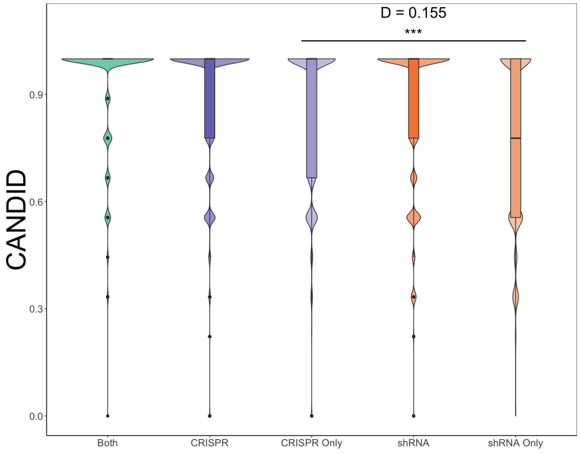


Figure S3

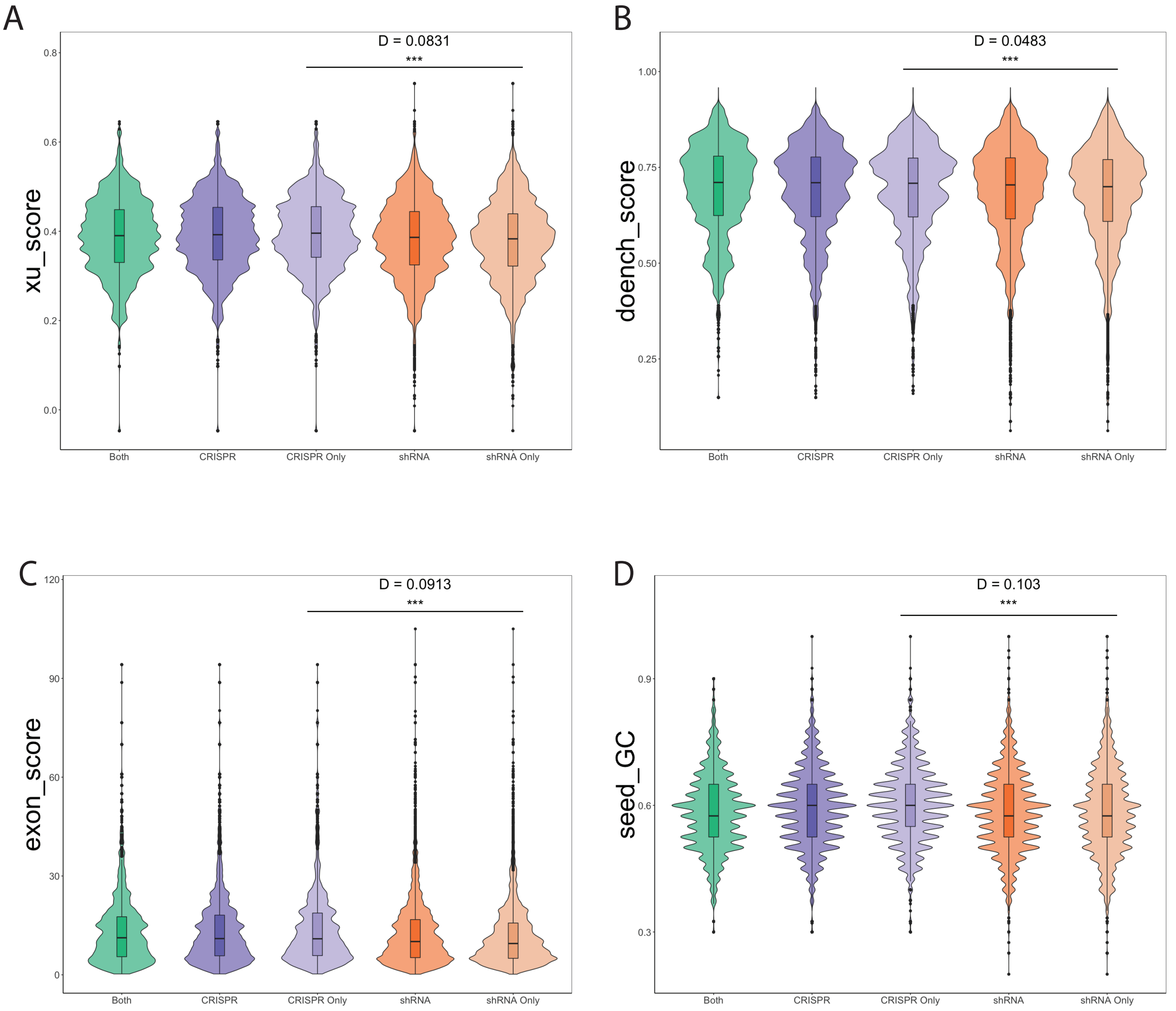

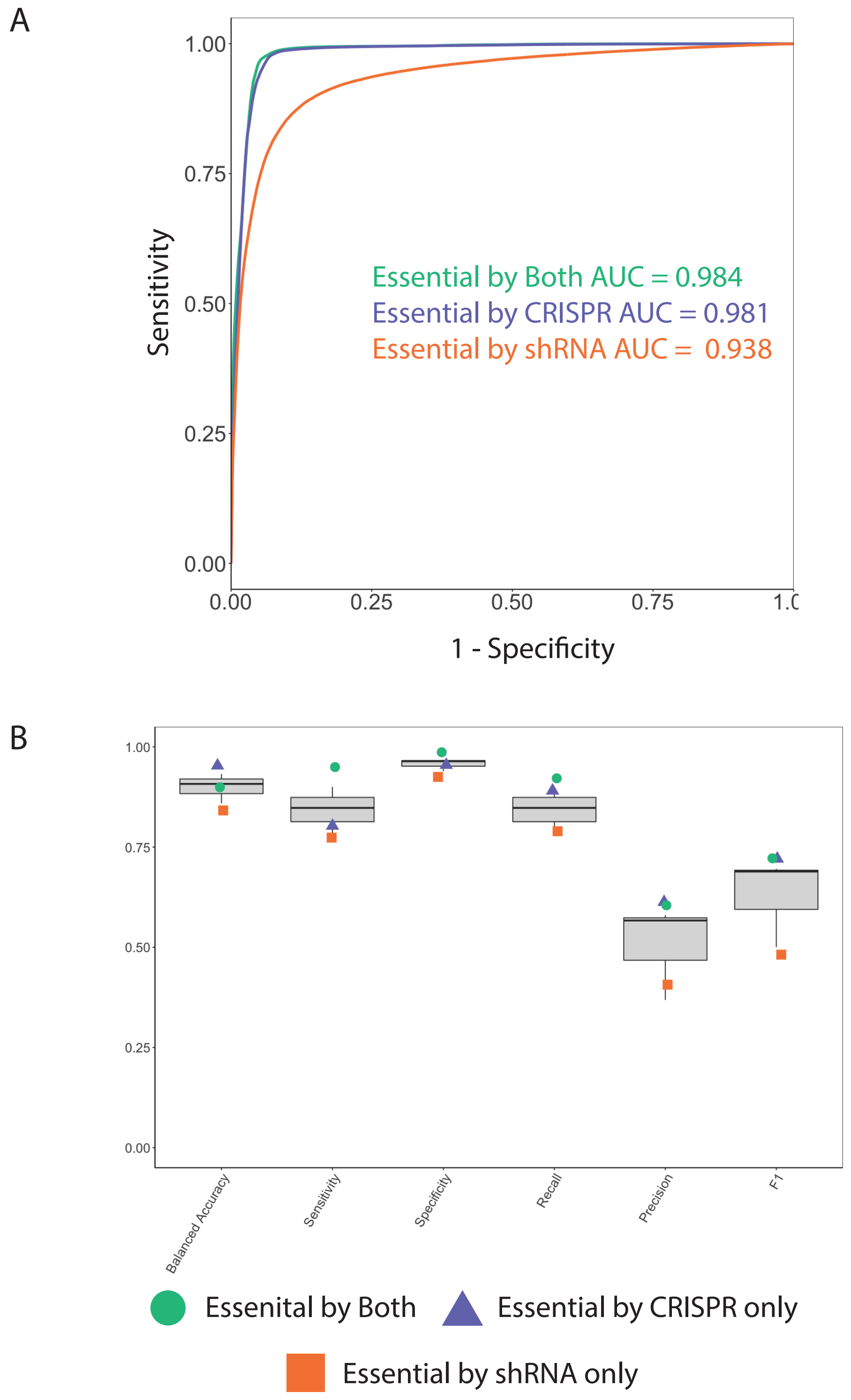

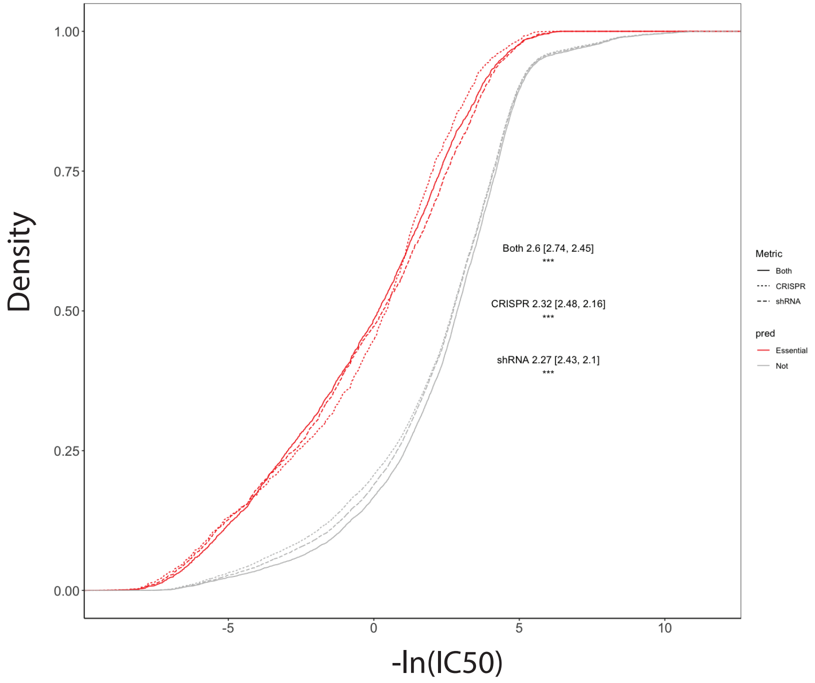

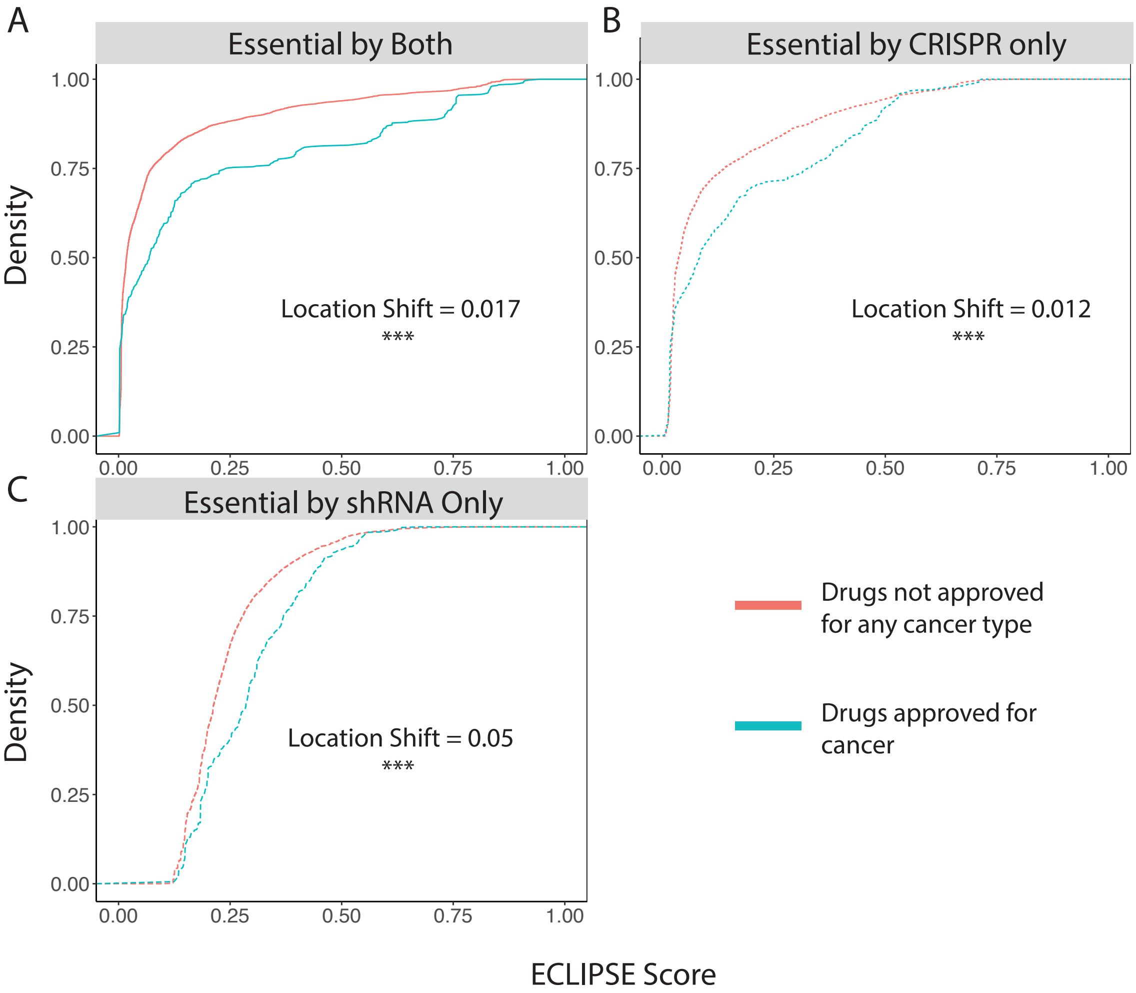
